# The genetics of eating behaviors: research in the age of COVID-19

**DOI:** 10.1101/2021.09.03.458854

**Authors:** Mackenzie E. Hannum, Cailu Lin, Katherine Bell, Aurora Toskala, Riley Koch, Tharaka Galaniha, Alissa Nolden, Danielle R Reed, Paule Joseph

**Author notes:** Corresponding Please address correspondence to: Danielle R Reed, PhD, Monell Chemical Senses Center, 35 Market St, Philadelphia PA 19104. equal contributions (Mackenzie E. Hannum, Cailu Lin).

## Abstract

How much pleasure we take in eating is more than just how much we enjoy the taste of food. Food involvement – the amount of time we spend on food beyond the immediate act of eating and tasting – is key to the human food experience. We took a biological approach to test whether food-related behaviors, together capturing food involvement, have genetic components and are partly due to inherited variation. We collected data via an internet survey from a genetically informative sample of 419 adult twins (114 monozygotic twin pairs, 31 dizygotic twin pairs, and 129 singletons). Because we conducted this research during the pandemic, we also ascertained how many participants had experienced COVID-19-associated loss of taste and smell. Since these respondents had previously participated in research in person, we measured their level of engagement to evaluate the quality of their online responses. Additive genetics explained 16-44% of the variation in some measures of food involvement, most prominently various aspects of cooking, suggesting some features of the human food experience may be inborn. Other features reflected shared (early) environment, captured by respondents’ twin status. About 6% of participants had a history of COVID-19 infection, many with transitory taste and smell loss, but all but one had recovered before the survey. Overall, these results suggest that people may have inborn as well as learned variations in their involvement with food. We also learned to adapt to research during a pandemic by considering COVID-19 status and measuring engagement in online studies of human eating behavior.

## 1. Introduction

The pleasure and perils of food start not on the tongue but, rather, well before the meal. Human eating behavior is more than the act of tasting, chewing, and swallowing food in the moment of eating – it also involves many preparatory and subsequent behaviors (Bell & Marshall, 2003), such as planning a meal, food shopping, cooking, and cleanup afterward. Drawing on studies demonstrating genetic effects on food preferences (Reed, Bachmanov, Beauchamp, Tordoff, & Price, 1997) and taste perception (Reed, Tanaka, & McDaniel, 2006), we took a biological approach to ask whether aspects of these other food behaviors have a genetic component. We tested this hypothesis using a genetically informative sample of adult human twins who answered questions about their own involvement with food via an online survey. These twins had participated in prior research projects about the sense of taste and smell (Knaapila, et al., 2012; Lin, et al., 2020; Wise, Hansen, Reed, & Breslin, 2007).

This research was conducted during the summer of 2020, when most research laboratories were closed due to the COVID-19 pandemic. Therefore, by necessity, we conducted the survey remotely. We also considered whether participants might have been ill or were currently ill with COVID-19 and how this might affect their responses. We knew at that time that taste and smell loss were cardinal features of COVID-19 illness (Parma, et al., 2020), more predictive of infection than other more general symptoms like fever (Gerkin, et al., 2021; Menni, et al., 2020), and that for some people the sensory loss lingers (Boscolo-Rizzo, et al., 2021). Therefore, it was reasonable to suspect that COVID-19 status, including the current state of a person’s ability to taste or smell, might affect responses to questions about food involvement (Hannum & Reed, 2021; Weir, Reed, Pepino, Veldhuizen, & Hayes, in review).

We were also concerned that participants who had participated in past research in a face-to-face setting, with supervision from investigators, might be less facile with remote research procedures. To address this concern, we included questions about engagement, drawing on a newly developed engagement scale, the Engagement Questionnaire (EQ), to discern the qualities of their responses: how active they were in thinking about the question, how much importance they attributed to the process, and how much they enjoyed taking the survey (Hannum & Simons, 2020). We could thus interpret participants’ responses about their food involvement by considering both their current or past taste and smell loss with COVID-19 and how well they adapted to the remote testing procedures.

## 2. Methods

### 2.1 Participants

We previously conducted research with human participants as part of the Twins Days Festival held annually in Twinsburg, Ohio (Knaapila, et al., 2012; Lin, et al., 2020; Wise, et al., 2007). As part of this research, the twins provided their contact information, including email addresses to be recontacted for future studies. Protocols compiled with the Declaration of Helsinki and were approved by the University of Pennsylvania Institutional Review Board (Protocol #843798). For data collection, we used this contact information to invite each twin to complete on online survey (described below) via REDCap, an electronic data capture tool used often in biomedical research (Harris, et al., 2009). The survey invitations were sent in waves starting September 3, 2020, and ending June 5, 2021; prospective participants who did not respond to the first message were sent an email reminder. All adult twins with internet access who could be reached by email were eligible to participate. All subjects provided informed consent to the research before starting the online survey. We collected demographic data from each participant, including their sex, age, race, and whether they were a current or former smoker or had never smoked in their lifetime. Participants received compensation for their time spent completing the survey.

### 2.2 Zygosity status

All twins surveyed had been previously genotyped as part of other research projects [4-6], and with appropriate consent, we used these genotypes to establish zygosity as monozygotic (MZ) or dizygotic (DZ). To establish zygosity, we relied on four methods. The twins self-reported on their zygosity, we took facial photographs which were rated for zygosity by two independent investigators, we typed all genomic DNA samples with a small panel of taste-related DNA markers, and for cases in which there was any uncertainty about the zygosity status using the first three methods, we genotyped the genomic DNA with the OmniChip from the Human OmniExpressExome-8v1.2 from Illumina Inc. (USA). In total, we genotyped 154 samples with the Omni-Chip and the remaining 265 samples using the first three methods.

### 2.3 Food involvement

Participants were asked to complete the Food Involvement Survey [1] (see **Table 1**), rating each item on a 7-point scale (1 = disagree strongly; 7 = agree strongly), as recommended in the original reference. The overall food involvement score has a theoretical range between 12 and 84, with higher numbers indicating more food involvement. The scale has two subscales: (1) presentation of the table and disposal of food and (2) preparation and eating of the food itself, in which certain items are summed into a final subscale score (see Table 1 for specific items).

**Table 1.** Food involvement questions and descriptive statistics

| Item | Trait | *n* | Question | Median | IQR |
| --- | --- | --- | --- | --- | --- |
| All | Total food involvement score<sup>a</sup> | 406 | All items scored and summed (range: 12-84) | 63 | 55-69 |
| Sub | Set and Disposal | 412 | Items 6, 11, 12 summed (range: 3 to 21) | 16 | 14-18 |
| Sub | Preparation/Eating | 408 | All other items summed (range: 9 to 63) | 47 | 41-52 |
|  |  |  | Individual items (range: 1-7) |  |  |
| 1* | Thinking about food | 413 | I don't think much about food each day | 2 | 2-4 |
| 2* | Cooking | 412 | Cooking or barbequing is not much fun | 2 | 2-4 |
| 3 | Talking about food | 413 | Talking about what I ate is something I like to do | 5 | 4-6 |
| 4* | Choice importance | 410 | My food choices are not very important | 2 | 2-4 |
| 5 | Travel | 413 | When I travel, one of the things I anticipate most is eating food there | 6 | 5-7 |
| 6 | Cleanup | 413 | I do most or all the cleanup after eating | 6 | 4-6 |
| 7 | Enjoy cooking | 411 | I enjoy cooking for others and myself | 5 | 4-6 |
| 8* | Eating out | 413 | When I eat out, I don't think or talk much about how the food tastes | 2 | 2-3 |
| 9* | Chopping | 413 | I do not like to mix or chop food | 2 | 2-4 |
| 10 | Food shopping | 413 | I do most or all of my own food shopping | 6 | 4-7 |
| 11* | Dishes | 412 | I do not wash dishes or clean the table | 1 | 1-2 |
| 12 | Setting table | 412 | I care whether or not a table is nicely set | 4 | 3-5 |

IQR=interquartile range. Some questions are paraphrased for brevity. Items are rated on a 7-point scale (1 = disagree strongly; 7 = agree strongly) and ordered by as they appeared in the original paper describing the Food Involvement Scale.
<sup>a</sup> The total food involvement score was generated by reversing the scores for items with an asterisk (\*) and then summing all scores.

### 2.4 Engagement

We measured engagement as the degree to which participants enjoyed, paid attention to, and made their best effort to respond to the questions in the survey. Our past research with these participants had been conducted in person, in an environment where they received face-to-face and individual instruction and supervision. Thus, in our view engagement was especially important to quantify for this internet survey, necessitated by our inability to conduct in-person research during the COVID-19 pandemic. We chose the newly developed Engagement Questionnaire (EQ) because it was developed specifically for sensory research and because of the expertise of the co-authors, one of whom developed it (Hannum & Simons, 2020). This tool as three subscales: active involvement (how vigorously they applied themselves to the task), purposeful intent (how they evaluated the importance or value of the survey), and affective value (how much they enjoyed the survey experience). We computed the average engagement scores from the subscales as described in the original report (Hannum & Simons, 2020).

### 2.5 COVID-19

This research was conducted during the COVID-19 pandemic (September 3, 2020-June 5, 2021). Because cardinal symptoms of COVID-19 are loss of taste, smell, and chemesthesis (e.g., the burn of chili pepper or the cool of menthol) (Parma, et al., 2020), it was important to establish whether participants had an active infection, or had been sick and lost their sense of taste or smell, because these senses affect food enjoyment (Hannum & Reed, 2021; Reed, et al., 2020). We therefore evaluated COVID-19 history using modified survey questions originally designed by members of the Global Consortium for Chemosensory Research (Parma, et al., 2020). We collected other measures for projects unrelated to the hypothesis tested here, and these measures are not reported.

### 2.6 Data analysis

#### 2.6.1 Data cleaning and descriptive analyses

To analyze the resulting data, we removed implausible age data, which we defined as having an age of >120 or <18 years (e.g., they provided the current date as their birthday). We next performed descriptive statistics on the food involvement total score, subscale scores, and individual item scores; engagement scores, subscale scores, and individual item scores; and COVID-19 measures, reporting the data as medians and interquartile ranges. Additionally, we calculated coefficient alpha (α) (Cronbach, 1951) to determine the internal reliability and consistency of each instrument, e.g., the Food Involvement Scale and Engagement Questionnaire).

#### 2.6.2 Covariate determination

Our goal was to establish the heritability of the food involvement traits, but first we needed to establish the appropriate covariates. To that end, the data from all participants were included in a linear regression to establish the influence of covariates on food involvement total score and individual items using age, sex, race, COVID-19 status, past chemosensory loss with COVID-19 (coded yes or no), and engagement subscale scores as dependent variables (Carlin, Gurrin, Sterne, Morley, & Dwyer, 2005). The participants who reported having had COVID-19 and past chemosensory loss did not differ in any food involvement scores compared to those without COVID-19 (**S1 Table**). Therefore, we dropped those COVID-19 factors from the model and reconducted the analysis. Sex, age, and engagement subscale scores were the most influential covariates (p<0.05) for at least some of the individual items on the Food Involvement Scale and were therefore included as covariates in the heritability analysis below. See **S2 Table** for details.

#### 2.6.3 Heritability analyses

Heritability was calculated using data from all 419 twin participants (114 MZ twin pairs (*N*=228 participants) and 31 DZ twin pairs (*N*=62), 129 singletons), using structural equation methods (Posthuma, et al., 2012). In this method, quantitative variation is parsed into variance attributable to additive genetic, shared environmental, and unique environmental effects using the *mets* package (version 1281) of R statistical software (Holst, Scheike, & Hjelmborg, 2016; Scheike, Holst, & Hjelmborg, 2014).

## 3. Results

### 3.1 Participants

We invited 1,742 twins to participate, and 434 (24.9%) responded by clicking the website link and attempting to complete the survey. We removed from the downstream data analysis anyone who did not consent (*N*=3), and those who provided implausible ages (*N*=12). After these data-cleaning steps, in total there were 419 participants (145 twin pairs (*N*=290) and 129 singletons). The participants were mainly female nonsmokers of European descent, with a median (IQR) age of 33 (29-46) years (**Table 2**). While 27 participants indicated they had COVID-19 at some point prior to the survey, and more than half of those had experienced taste and/or smell loss when they were ill, only one person indicated problems with current ability to taste or smell. Regardless, food involvement scores did not differ significantly between those infected with SARS-CoV-2 and experienced chemosensory loss and those that did not (**S1 Table**).

**Table 2:** Participant characteristics

| Characteristic | N | % |
| --- | --- | --- |
| Total participants | 419 | 100 |
| Currently able to taste/smell | 418 | 99.8 |
| Age [median (IQR), years] | 33 (29-46) |  |
| Sex |  |  |
| Female | 346 | 82.6 |
| Male | 72 | 17.2 |
| Prefer not to say | 1 | 0.2 |
| Ancestry |  |  |
| European | 368 | 87.6 |
| African | 22 | 5.3 |
| Asian | 12 | 2.9 |
| Other | 17 | 4.1 |
| Smoking |  |  |
| Never | 329 | 78.5 |
| Ever | 77 | 18.4 |
| No response | 13 | 3.1 |
| Twin status |  |  |
| MZ | 114 | 54.4 |
| DZ | 31 | 14.8 |
| Singleton | 129 | 30.8 |
| COVID-19 infection history | 27 | 6.4 |
| Taste change | 19 | 4.5 |
| Smell change | 17 | 4.1 |

MZ=monozygotic, DZ=dizygotic.

### 3.2 Food involvement scores

Median scores and their IQRs for total food involvement and the individual items are provided in **Table 1**. Items are coded on a 7-point scale (1 = disagree strongly; 7 = agree strongly). The median scores were 5 or 6 for most positive items (e.g., *I enjoy cooking*) and 2 for most negative items (e.g., *Cooking or barbequing is not much fun*). The most variable trait based on the IQR was food shopping (*I do most or all of my food shopping*). The median total food involvement score, calculated by reversing the scores for the negative items and then summing scores for all items, was 63, with an IQR of 55–69. Overall, the Food Involvement Scale had a Cronbach’s α of 0.77 (95% CI: 0.73-0.80), demonstrating acceptable internal reliability.

### 3.3 Effects of engagement

Participants’ engagement scores showed that most had made their best effort (active involvement, median = 5.7 and IQR = 4.7-6.0), considered the research important (purposeful intent; median = 5.5 and IQR = 5.0-6.0), and enjoyed answering the questions (affective value; median =4.7 and IQR = 4.0-5.3). With a Cronbach’s α of 0.84 (95% CI: 0.82-0.87), the Engagement Questionnaire has internal consistency. Overall, participant’s level of engagement was significantly related to their level of reported food involvement (Table 3). Specifically, whether participants thought the research and their role as a participant were important (captured via the purposeful intent subscale) had the strongest effect on their reported food involvement score and sub-scale score (coefficients >1, Table 3). Whether subjects were actively involved in the task did not influence their reported level of involvement with set and disposal procedures of consuming food. Including questions about engagement allowed us to assess the participant’s level of engagement in the online task, and control for lack of attention, increasing the accuracy of the heritability estimates.

**Table 3:** Engagement scale significantly correlated with food involvement scores (Coefficient [95%CI]).

| Items | n | Active involvement | Purposeful intent | Affective value |
| --- | --- | --- | --- | --- |
| Food involvement score | 391 | **\*\*\*1.76 [0.89 - 2.63]** | **\*\*\*3.93 [2.52 - 5.35]** | **\*\*\*2.61 [1.49 - 3.74]** |
| Preparation and eating | 393 | **\*\*\*1.52 [0.74 - 2.30]** | **\*\*\*2.78 [1.60 - 3.95]** | **\*\*\*1.81 [0.88 - 2.74]** |
| Set and disposal | 395 | 0.24 [-0.03 - 0.50] | **\*\*\*1.14 [0.67 - 1.60]** | **\*\*\*0.77 [0.43 - 1.12]** |
| Thinking about food | 397 | -0.14 [-0.31 - 0.02] | 0.06 [-0.17 - 0.29] | 0.06 [-0.12 - 0.25] |
| Cooking | 396 | **\*\* -0.23 [-0.37 - -0.09]** | **\*\*\* -0.51 [-0.71 - -0.3]** | **\*\*\* -0.33 [-0.51 - -0.15]** |
| Talking about food | 397 | 0.03 [-0.13 - 0.19] | **\*\*0.3 [0.07 - 0.52]** | **\*\*\*0.29 [0.12 - 0.46]** |
| Choice importance | 396 | **\*\*\* -0.33 [-0.46 - -0.2]** | **\*\*\* -0.42 [-0.63 - -0.22]** | **\*\* -0.23 [-0.39 - -0.07]** |
| Travel | 397 | -0.05 [-0.19 - 0.09] | 0.01 [-0.20 - 0.22] | 0.02 [-0.14 - 0.18] |
| Cleanup | 397 | **\*0.14 [0.02 - 0.25]** | **\*\*\*0.39 [0.20 - 0.58]** | **\*\*\*0.28 [0.13 - 0.42]** |
| Enjoy cooking | 395 | **\*\*\*0.29 [0.14 - 0.45]** | **\*\*\*0.61 [0.40 - 0.83]** | **\*\*\*0.41 [0.22 - 0.6]** |
| Eating out | 397 | -0.07 [-0.19 - 0.05] | -0.18 [-0.36 - 0.00] | -0.08 [-0.24 - 0.07] |
| Chopping | 397 | **\*\* -0.21 [-0.35 - -0.08]** | **\*\* -0.32 [-0.52 - -0.13]** | **\* -0.2 [-0.37 - -0.02]** |
| Food shopping | 397 | **\*0.19 [0.04 - 0.34]** | **\*\*\*0.42 [0.19 - 0.64]** | **\*0.22 [0.04 - 0.41]** |
| Dishes | 396 | **\*\* -0.15 [-0.26 - -0.04]** | **\*\*\* -0.35 [-0.55 - -0.15]** | **\* -0.19 [-0.34 - -0.03]** |
| Setting table | 396 | -0.04 [-0.19 - 0.10] | **\*\*\*0.38 [0.16 - 0.60]** | **\*\*0.3 [0.10 - 0.49]** |

Note: Bolded values are significant; \*p<0.05, \*\*p<0.01, \*\*\*p<0.001; Coefficient [95% CI] was determined in the twin regression model for each single engagement sub-scale separately.

### 3.4 Heritability of the food involvement scores

About 40% of individual differences in food involvement scores was explained by additive genetic variance (additive genetics; a^2^ = 0.39; 95% CI = 0.24–0.55, *P*<0.001). However, although food involvement could be considered a single entity, each item from the scale contributed to this result in a different way (**Figure 1** and **S3 Table**). The traits for cooking (“*I enjoy cooking for myself and others*”; a^2^ = 0.40; 95% CI = 0.25–0.56, *P*<0.001) and especially chopping (“*I do not like to mix or chop food*”; a^2^ = 0.44; 95% CI = 0.30–0.58, *P*<0.001) were the most highly heritable as measured by additive genetics, which points to a heritable component for food preparation. We observed a similar but weaker effect for the trait ‘dishes’ (“*I do not wash dishes or clean the table*”; a^2^=0.17; 95% CI =0.01 - 0.32, p<0.001].

**Figure 1.**
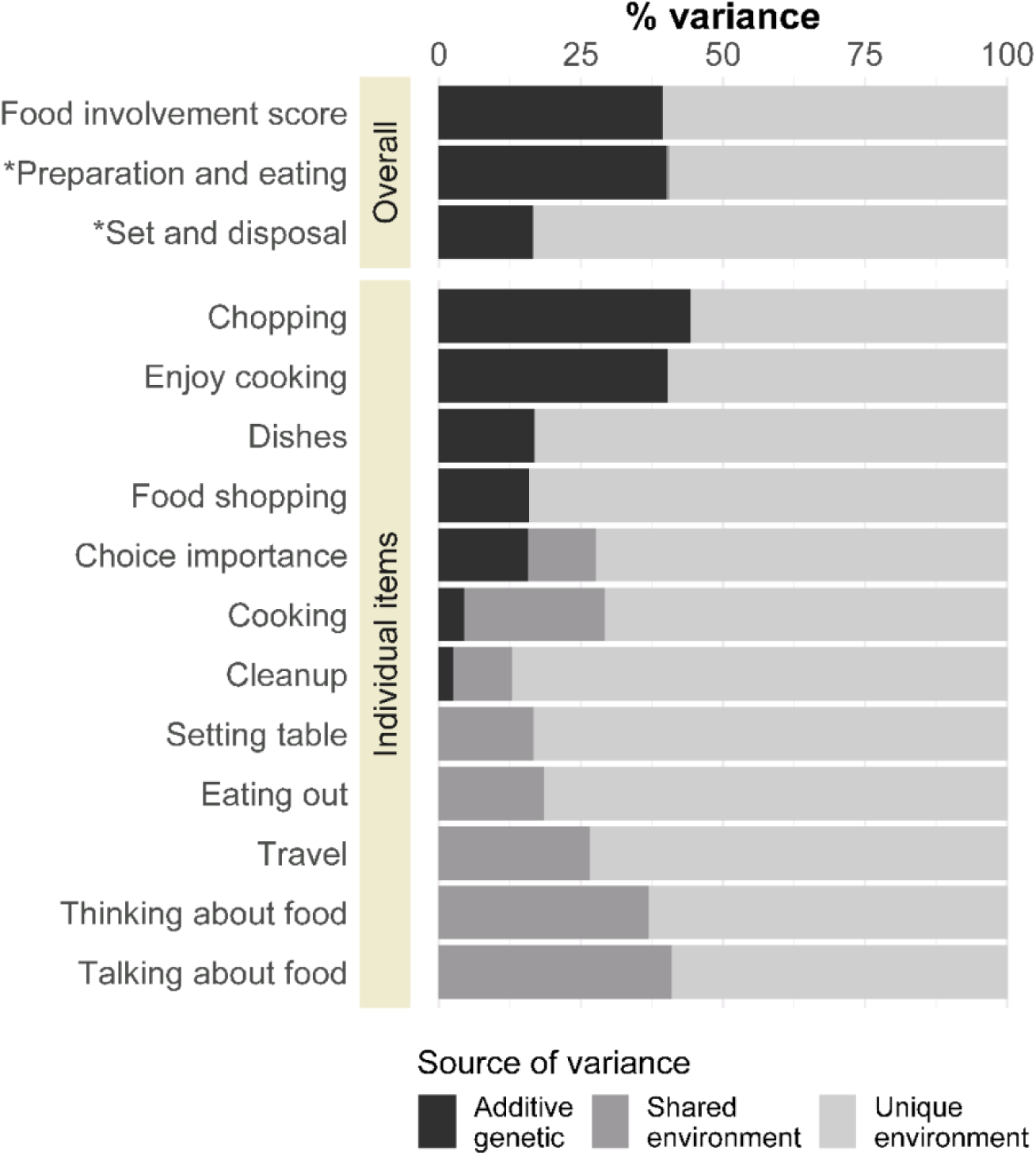
Variance in food involvement, by additive genetic, shared environment, and unique factors. Food involvement score refers to the total score, computed as described in section 1.3. Scores for the two subscales (labeled with *) and the separate items are below (for fuller description, see Table 1), ordered from more to less additive genetic variance.

The effects of a shared environment, for which twins are the most similar regardless of zygosity, are captured by questions assessing thinking (c^2^ = 0.37; 95% CI = 0.22–0.52, *P*<0.001) and talking about food (c^2^ = 0.41; 95% CI = 0.27–0.55, *P*<0.001) and interest in food during travel (c^2^ = 0.26; 95% CI = 0.12–0.41, *P*<0.001). The trait most influence by the unique environment unshared among twins was cleanup (“*I do most or all the cleanup after eating*”; e^2^ = 0.87; 95% CI = 0.70–1.04, *P*<0.001). Drawing on the subscales to help summarize these results, food preparation is more heritable (Preparation and Eating subscale, a^2^ = 0.40; 95% CI = 0.38–1.18, P<0.001) and cleanup afterward is generally less so (Set and Disposal subscale, a^2^ = 0.17; 95% CI = -0.01–0.34, *P*<0.001).

## 4. Discussion

While much research into the biology of human food behavior focuses on the moment of eating – how much is eaten and the types of food chosen, e.g.,(Grimm & Steinle, 2011; Pallister, Spector, & Menni, 2014) – or even being fearful of new foods (Knaapila, et al., 2007), other types of behavior are also important parts of the total food experience, including the selection and preparation of food, ruminating about food, looking forward to eating in new places, and the cleanup afterward (Bell & Marshall, 2003). The results of our online survey of twins show that there is a genetic determinant to at least some aspects of these food behaviors, with cooking and food preparation the most heritable (i.e., genetically identical twins were more similar in these types of behavior than were fraternal twins).

However, while cooking, and especially the chopping and mixing of food, had a heritability component, other types of food behaviors were affected more by shared common environment, and twins were thus similar regardless of their degree of genetic relatedness (i.e., genetically identical twins were as similar in these types of behavior than dizygotic twins). The most striking examples of this effect of shared common environment are the scores for talking and thinking about food, with more than 40% of the variance in these traits accounted for being raised in a shared household.

Some aspects of food and eating were unaffected either by genetics or by shared common environment and thus appeared to be determined by each person’s own experience. The most prominent examples of this were the results for setting the table and cleaning up afterward, which were unrelated to shared experience or genetics between the twins.

These results suggest that the proclivity for cooking may be more determined by genetic variation, whereas the centrality of food in the family – how much importance is attached to it – may be more learned in the family home. Taken together, it appears that love (or dislike) of cooking may be difficult to modify but that the centrality of food (e.g., as topic of conversation) may be influenced by the family environment and that the willingness to set up and clean-up is entirely amenable to change (e.g., through learning or instruction, or by the encouragement of certain behaviors).

This research was conducted in unusual circumstances because of the pandemic. We learned from this research that, while a handful of the participants had COVID-19 at some point immediately before the survey, only one person still had problems with taste and smell at the time of their survey response. We saw no differences in food involvement scores based on whether the participants had past or active COVID-19 infection, and it is encouraging that people recovered with little or no effect in their food behaviors. While only one person reported problems with taste and smell, studies of recovery suggest that 10% or more of people have sustained taste and smell loss after many months (Boscolo-Rizzo, et al., 2021). Though not tested presently, we hypothesize that people with long-haul COVID-19 symptoms and sustained taste and smell loss might have much less food enjoyment (Boesveldt, et al., 2017; Temmel, et al., 2002), however, its resultant effect on food involvement has yet to be determined. Since food involvement is thought to be a stable individual characteristic (Bell & Marshall, 2003), it would be of interest to explore how sudden and *sustained* loss of taste and/or smell might affect someone’s overall level of food involvement. Regardless, questions about COVID-19 and its effect on taste and smell should be included in human food research in future, as a basic demographic question like age or sex.

We quantified engagement because we were concerned about the abrupt change of testing procedures from in person to remote, owing to the laboratory lockdowns during COVID-19. This decision proved to be more fruitful than we anticipated. Not only did participants’ scores reassure us that most had made their best effort, considered the research important, and enjoyed answering the questions, but we found that aspects of engagement – whether participants thought the research and their role as a participant were important – were strongly related to participants’ food involvement scores. An aspect of individual food involvement encompasses a higher level of cognitive interest in a task related to food, such as explaining the differences between products or discussing food in general (Bell & Marshall, 2003). Thus, participants who reported a higher level of food involvement additionally reported higher levels of purposeful intent during the survey suggesting they found value in discussing food-related topics. This supports the converging (e.g., similar) aspects between engagement and food involvement, an important aspect to using validated scales in research (DeVellis, 2016). In general, including questions about engagement allowed us to validate using a remote survey tool and increase the accuracy of the heritability estimates, similar to using age and sex as covariates.

This research has at least two limitations. First, because of our recruitment strategy – former attendees and research participants from an annual festival for twins – we had more genetically identical than fraternal twin pairs, which might reduce our power to detect additive genetic variance. Second, some twins responded but their co-twin did not. We attribute this situation to two factors (neither of which was in our immediate control): some contact information was outdated, so we could not reach the co-twin; and the vicissitudes of COVID-19 and the pandemic, with constant upheaval and changes, made it hard for some to prioritize participating in research. These singletons are not without value, because they provided information for estimating the effects of covariates and also about variation when computing heritability. However, we recognize that it would be more valuable from a genetics perspective to have all twin pairs in our sample rather than include singletons.

The heritable variations we identified in interest and liking for cooking and food preparation are not surprising, if we consider that cooking is often a recreational or leisure activity, like competitive or team sports, which are often related to heritable traits (e.g., (van der Zee, Helmer, Boomsma, Dolan, & de Geus, 2020)). However, this observation takes on new importance in the realm of health, because the willingness to cook and prepare food at home is related to better overall health (Wolfson & Bleich, 2015) and people who have high food involvement scores have a healthier diet (Barker, Lawrence, Woadden, Crozier, & Skinner, 2008; Marshall & Bell, 2004). Our results on the genetics of food involvement have implications for personalized nutrition and how much behavior change is likely to occur when people who do not like to cook at home are encouraged to do so. These genetic aspects on the enjoyment of cooking may constrain behavior change. However, more research is needed to explore this hypothesis.

In contrast, how much people are preoccupied with thinking and talking about food is highly affected by shared family environment, which suggests that the standards set in the childhood home will persist into adulthood. Some families are more focused on food as a hub of life, whereas others are less so, and level of parental emphasis on food may have effects into adulthood. We hasten to add that the questions on the Food Involvement Scale do not address pathological preoccupation with food, such as with eating disorders including anorexia and bulimia, which have a strongly heritable component (Thornton, Mazzeo, & Bulik, 2011). Rather, social discussion and interest in food are affected by shared environment, at least in this population of human twins.

Looking to the future, as the collective data from genetic studies bring larger and larger databases and more available genotyping information, we will learn about how inborn variation affects what people choose to eat (Cole, Florez, & Hirschhorn, 2020; May-Wilson, et al., 2021). Our hope is that, as the capacity for this type of research grows, behaviors such as those studied here, which are key parts of the full human experience of food, will be included in future research. Although ultimately human health is improved directly by what and how much people eat, these decisions are made as people shop, cook, linger over one meal, and anticipate the next.

## Figure Caption

**Figure 1**. Variance in food involvement, by additive genetic, shared environment, and unique factors. Food Involvement Scale refers to the total score, computed as described in section 1.3. Scores for the two subscales (labeled with *) and the separate items are below (for fuller description, see Table 1), ordered from more to less additive genetic variance.

## Abbreviations

EQ: Engagement Questionnaire
FIS: Food Involvement Score
MZ: monozygotic
DZ: dizygotic

## Acknowledgments

We thank Maija Aurora Greis, Allison Cox, Elizabeth Cole, Rongze Sun, Sarah Lipson, Sarah Marks, and Corrine Mansfield for their assistance with data management. We are grateful to the twins for their participation in a time of upheaval and uncertainty and to the Twins Days Festival organizers including Sandy Miller and Janine Bregitzer. We acknowledge the assistance of the following people in data collection: Laura Alarcon, Kelsey Alderman, Tiffany Weiss Aleman, Charles J. Arayata, Nuala Bobowski, Seth Brockman, Lauren Colquitt, Danielle Crawford, Laurel Doghramji, Jennifer Douglas, Fujiko Duke, Hillary Ellis, Emily Evans, Brad Fesi, Brian Gantick, Federica Genovese, Nicole Greenbaum, Rebecca James, Desmond Johnson, Amin Khosnevisan, Matthew Kirkey, Antti Knaapila, Marie Knoll, Daniel Lee, Deborah Lee, John Lees, Anna Lysenko, Ivy Maina, Alex Mangroo, Corrine Mansfield, Michael Marquis, Elliott McDowell, Michelle Murphy Niedziela, Kevin Redding, Jake Saunders, Lauren Shaw, Molly Stein, Alyssa Treff, Casey Trimmer, Tom Uleau, Calvin Alarcon Uleau and Ana Ulloa.

## Author Contributions

MEH designed the study, cleaned the data, assisted in the statistical analysis and contributed to the writing of the manuscript, CL performed the statistical analysis and contributed to the writing of the manuscript, KB, AT, and RK collected data and edited the manuscript, TG contributed to the design of the study, AN contributed to the design of the study, assisted in data collection and assisted in the writing of the manuscript. DRR assisted in the design of the study, assisted in data analysis, and wrote the manuscript. PJ designed the study and assisted in the writing of the manuscript.

## Ethical statement

The study was conducted according to the guidelines of the Declaration of Helsinki and approved by the University of Pennsylvania Institutional Review Board (Protocol #843798). The procedures performed were in accordance with the ethical standards of the committee. Electronic informed consent was obtained from all respondents before commencement of the study.

